# Activity-dependent synthesis of translational machinery through dynamic remodeling of ribonucleoprotein granules

**DOI:** 10.64898/2026.06.28.734671

**Authors:** Toshiharu Ichinose, Mai Kanno, Takashi Makino, Yuichi Shichino, Shintaro Iwasaki, Hiromu Tanimoto

## Abstract

Neuronal stimulation drives gene-expression programs supporting long-lasting changes, yet how depolarization reshapes post-transcriptional regulation remains poorly understood. Here, we combined optogenetic stimulation with neuron-specific Ribo-seq and RNA-seq in the *Drosophila* brain to resolve activity-evoked translational dynamics. Stimulation triggered two temporally ordered programs: a rapid wave to translate synapse remodeling factors and neuropeptides, followed by a dominant delayed program that selectively upregulated translation of protein synthesis machinery, including nearly all cytosolic ribosomal proteins (RPs). Transcripts in the late program are marked by 5’ terminal oligopyrimidine (TOP) motifs. TOP mRNAs accumulated in ribonucleoprotein granules at the basal state but are released upon stimulation. We identified La-related protein as a key regulator of the activity-dependent translation of TOP mRNA and memory consolidation. Comparative genomics indicated that TOP enrichment in RP mRNAs is specific to metazoans. These findings suggest that animals acquired activity-dependent TOP-mRNA control to couple neuronal stimulation with adaptation of translational capacity.

## Introduction

Neurons convert transient neuronal stimulation into lasting changes in synaptic strength, circuit function, and behavior. Activity-dependent gene-expression programs underlie these plastic changes. Seminal studies identified immediate early genes (IEGs) as rapid transcriptional targets of neuronal activation and established their broad roles in neuronal adaptation ^1–5^. More recently, transcriptome-wide approaches have systematically cataloged genes induced by defined stimulation paradigms ^6,7^ or by learning-related experiences ^8–10^. Nevertheless, transcriptional changes alone are insufficient to explain the full range of activity-dependent neuronal adaptations, which also require post-transcriptional regulation ^11,12^.

Translation offers a rapid and spatially flexible route to remodel the neuronal proteome, since it can utilize pre-existing mRNAs stored in subcellular compartments, including ribonucleoprotein (RNP) granules ^13,14^. A seminal study in hippocampal slices showed that neurotrophin-induced, long-lasting enhancement of synaptic transmission requires protein synthesis, even when neuropiles are isolated from the cell bodies ^15^. Similarly, repeated stimulation of motor neurons in *Drosophila* induces translation of pre-existing CaMKII mRNA ^16^. Behavioral studies further demonstrated that early long-term memory depends on *de novo* protein synthesis without new transcription, emphasizing learning-induced translational programs during memory consolidation ^17,18^. Consistent with these observations, studies have reported that persistent synaptic plasticity and memory formation involve multiple translational regulators ^19–24^. Nevertheless, activity-induced genome-wide translational remodeling in intact brains remains poorly defined.

Our recent study established an experimental strategy for *in vivo* cell-type resolved translational profiling by combining genetic tagging of ribosomes with ribosome profiling (Ribo-seq), which quantifies ribosome-protected mRNA footprints at nucleotide resolution ^25–27^. Here, we leverage neuron-specific ribosome profiling and optogenetic stimulation in the *Drosophila* brain to obtain a time-resolved transition of activity-evoked translation. We find that neuronal activity sequentially enhances translation of synapse-remodeling factors and, more prominently, the protein synthesis machinery itself. The latter program encompasses ribosomal proteins (RPs) and translation factors, containing 5’ terminal oligopyrimidine (TOP) motifs. We further show that TOP mRNAs are stored in RNP granules and released for translation in a Larp-dependent manner, that this regulation gates memory consolidation, and that selective TOP enrichment among RP mRNAs is a metazoan-specific feature. These findings reveal a metazoan-evolved mechanism by which neuronal activity controls translational capacity.

## Results

### Neuronal stimulation drives immediate early translation of specific genes

To investigate activity-dependent translatome dynamics in the *Drosophila* brain, we combined optogenetic stimulation and neuron-specific ribosome profiling. We expressed *channelrhodopsin-2-XXL* (*ChR2XXL*) ^28^ together with an epitope-tagged ribosomal protein, *RpL3-FLAG* (uL3 in universal nomenclature) ^26,29^ under the control of the pan-neuronal driver *nSyb-GAL4*. Following a three-second light stimulus, we isolated neuron-specific ribosome-bound mRNAs by immunoprecipitation followed by ribosome footprinting (Figure 1A). Rapid flash-freezing enabled us to monitor translational events at precisely defined time points after stimulation (2, 10, 30, 90, and 270 min). We validated neuron-specific ribosome profiling by confirming enrichment of neuronal marker genes (Figure S1A), depletion of markers from non-neuronal cell types (Figure S1A), robust three-nucleotide periodicity (Figure S1B), and highly consistent expression patterns with or without ChR2XXL expression (Figure S1C).

**Figure 1.**
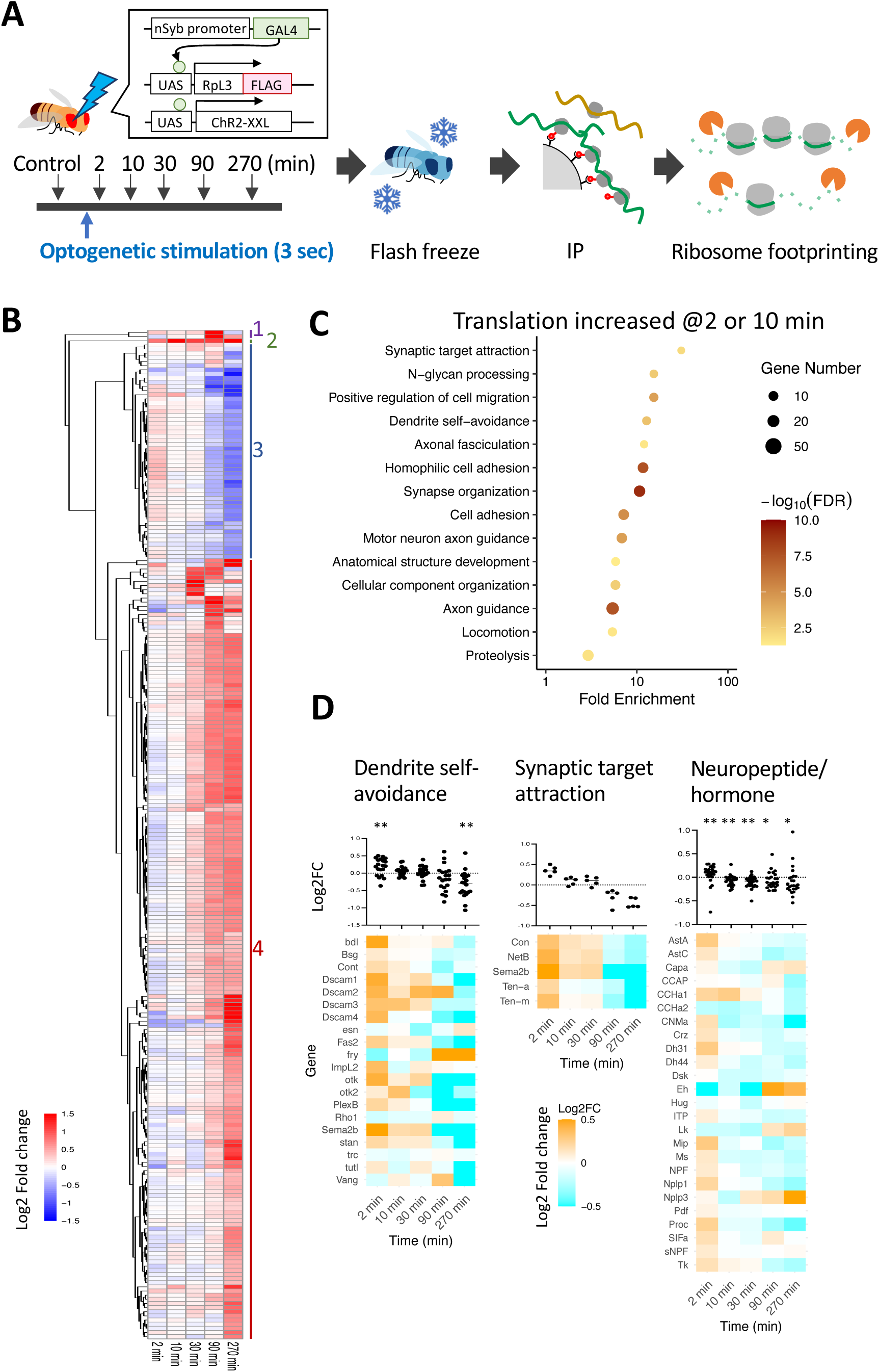
Immediate early translation after neuronal stimulation. **(A)** Experimental design for time-resolved neuron-specific ribosome profiling after optogenetic stimulation. Flies expressing ChR2-XXL and FLAG-tagged ribosomal protein L3 in neurons were stimulated for 3 sec, and heads were collected at the indicated time points (Control, 2, 10, 30, 90, and 270 min). Flies were flash-frozen with liquid nitrogen and the heads were collected. Following immunoprecipitation with anti-FLAG antibody, ribosome footprinting was performed. **(B)** Heatmap of transcripts showing significant increases or decreases in ribosome footprints on CDS (*FDR* < 0.05, DESeq2). Color indicates log2FC relative to control. Genes are hierarchically clustered by temporal pattern, and numbers on the right indicate cluster numbers. Gene names are shown in Figure S2. **(C)** GO enrichment analysis of transcripts with increased translation at early time points after stimulation (2 or 10 min, *P* < 0.05, DESeq2). Dot size indicates gene number, and color indicates enrichment significance (−log10[FDR]). **(D)** Heatmaps showing time-course translational changes of representative genes in functionally enriched categories, including “dendrite self-avoidance” (GO:0070593), “synaptic target attraction” (GO:0016200), and “neuropeptide hormone activity” (GO:0005184). Values are log2 fold changes relative to unstimulated control. Asterisks indicate statistical significance against 0 (* *P* < 0.05, ** *P* < 0.01, Wilcoxon signed-rank test).

We identified 243 genes with significant changes in ribosome occupancy (*FDR* < 0.05 in DESeq2 ^30^) at one or more time points after stimulation. Hierarchical clustering grouped these genes into four distinct clusters (Figure 1B). Cluster 3 included genes with rapid induction at 2 and 10 min followed by later repression, whereas cluster 4 showed delayed and sustained upregulation from 30 to 270 min, revealing two temporally distinct translational programs (Figure 1B). To characterize the early response, we analyzed genes with increased ribosome occupancy at 2 or 10 min relative to the unstimulated controls (*P* < 0.05). Gene Ontology (GO) analysis revealed significant enrichment of “dendrite self-avoidance,” “synapse organization,” and “axon guidance” (Figure 1C), suggesting rapid synthesis of proteins involved in synapse and circuit remodeling. Consistently, guidance and adhesion molecules such as *Sema2b*, *Ten-a*, *Ten-m*, and *Dscam* showed sharp and transient translational induction within 10 min (Figure 1D).

The early program also encompassed neuropeptides, including *AstA*, *Mip*, and *Dh31*: 21 of 25 neuropeptide genes exhibited higher ribosome occupancy than unstimulated controls, with peak induction at 2 min (Figure 1D). This rapid upregulation may represent immediate feedback replenishment after activity-triggered neuropeptide release.

### Systematic translational upregulation of genes involved in protein synthesis

In contrast to the early translational response, we found a dominant late program characterized by coordinated upregulation beginning 30 min after stimulation (cluster 4). Strikingly, approximately 40% of genes in this translationally enhanced cluster encoded ribosomal components, representing 78 of 80 cytosolic RPs (97.5%; Figure 2A). Mitochondrial RPs did not show significant upregulation (Figure S5), indicating selective and systematic translational enhancement of cytosolic ribosome components. GO analysis further revealed enrichment for “cytosolic translation,” “ribosome biogenesis,” “translational initiation/elongation,” and “protein folding” (Figures 2B and S3). Consistently, the late program included ribosome-assembly regulators (*Ip259*, *Nop5, Nop56, Nop60B,* and *sta*), five initiation factors and four elongation factors, poly(A)-binding protein (*pAbp*), six heat-shock proteins, and the CCT chaperonin complex (Figures S3 and S4). Thus, the largest activity-dependent translational program enhances the protein synthesis machinery itself.

**Figure 2.**
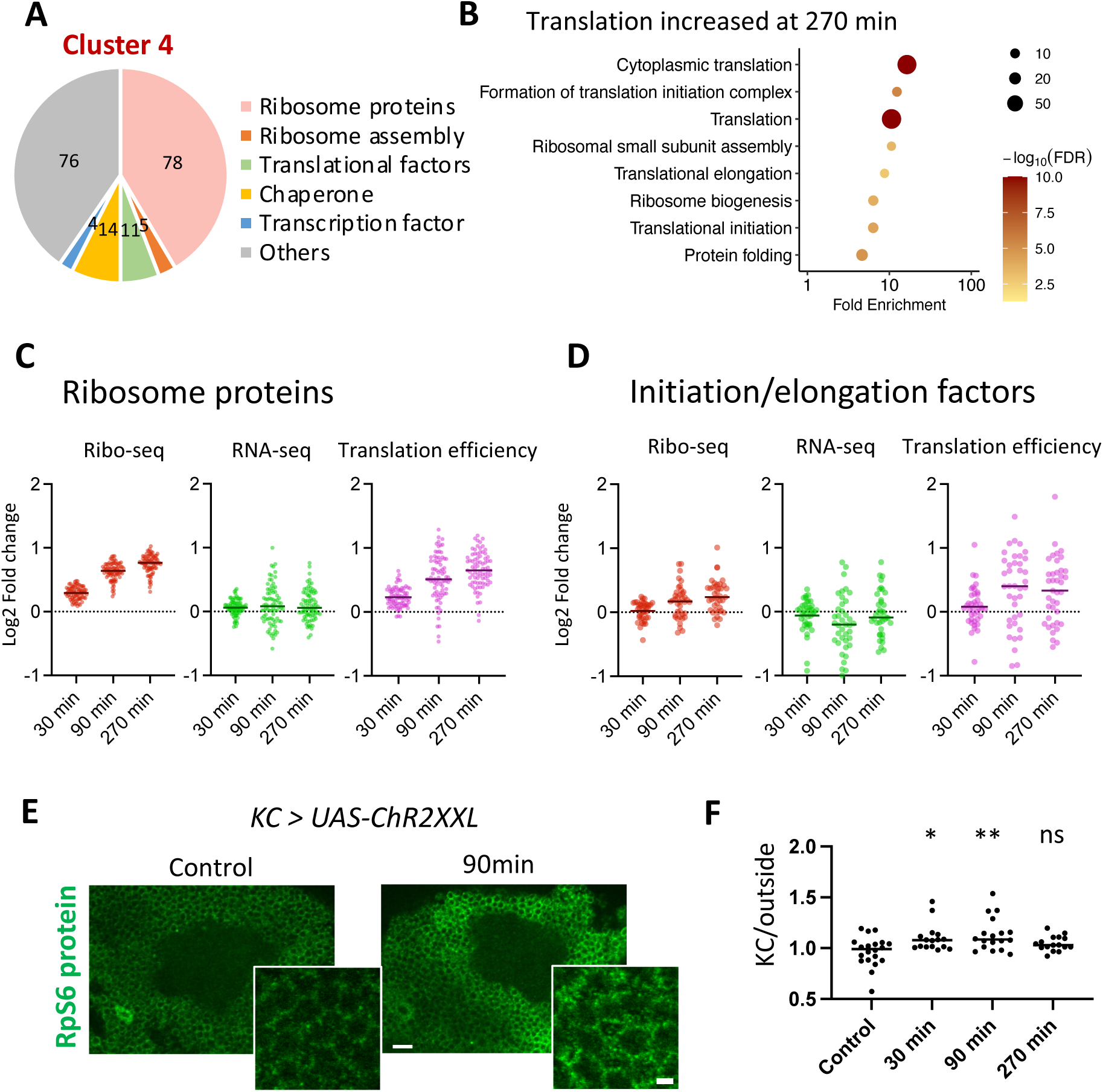
Translation machinery is synthesized through post-transcriptional regulation. **(A)** Functional classification of genes in cluster 4, corresponding to dominant translational induction, shows enrichment of ribosomal proteins, ribosome assembly factors, translational factors, chaperones, and transcription factors. Numbers indicate gene counts in each category (188 total genes). **(B)** GO enrichment analysis of genes with increased translation at 270 min after stimulation. Dot size indicates gene number, and color indicates -log10(FDR). **(C)** Time-course changes in ribosomal protein genes in Ribo-seq, RNA-seq, and translational efficiency (TE), showing translational enhancement with minimal mRNA induction. **(D)** Time-course changes in translation initiation and elongation factor genes in Ribo-seq, RNA-seq, and TE. **(E)** Representative immunohistochemistry images of RpS6 protein in KCs with or without optogenetic stimulation. *ChR2XXL* was expressed using *tubP-GAL80^ts^;R13F02-GAL4*, and flies were optogenetically activated for 30 sec (see Methods for details). Insets show magnified regions. Scale bars, 10 µm (main) and 2 µm (inset). **(F)** Quantification of RpS6 signal, calculated as the ratio of signal intensity in KCs to that outside KCs at the indicated time points after stimulation. Each dot represents one animal, and lines indicate medians. Asterisks denote significance; ns, *P* > 0.05, *: *P* < 0.05, **: *P* < 0.01, Dunn’s multiple-comparison test versus control.

To compare these translatome changes with transcriptional responses, we performed neuron-specific RNA-seq (TRAP-seq) after the same optogenetic stimulation paradigm at 30, 90, and 270 min. Previously reported activity-regulated genes ^7^ were similarly upregulated in our transcriptome dataset (Figure S6). By contrast, mRNA abundance of RPs and initiation or elongation factors underwent limited changes (Figures 2C and 2D), indicating that neuronal activity primarily regulates these genes at the level of translation. We validated this delayed program at the protein level by immunohistochemistry for ribosomal protein S6 (RpS6 or eS6) upon optogenetic stimulation of mushroom body intrinsic neurons, the Kenyon cells (KCs; Figure 2E and 2F). Together, these data reveal that neuronal activity upregulates the protein synthesis machinery via translation rather than transcription.

Beyond the protein synthesis machinery, we observed increased translation of cytoskeletal proteins (*Act5C*, *αTub56D*, and *αTub84B*) at later time points (Figures S2 and S3), suggesting cytoskeletal remodeling after the early synaptic responses. Oxidative stress response genes (*GstE4*, *Trxr-1*, *Ldsdh1*, and *Fmo-2*) were also upregulated (Figure S4), consistent with the increased energetic and redox demands associated with these processes.

### The TOP mRNA motif underlies activity-dependent translation

Transcripts encoding cytosolic RPs contain a 5’ terminal oligopyrimidine (TOP) motif ^31,32^. Motif-enrichment analysis ^33^ revealed a significant enrichment of pyrimidines (C and U) immediately downstream of the 5’ cap among cluster 4 genes (Figures 3A and S7).

**Figure 3.**
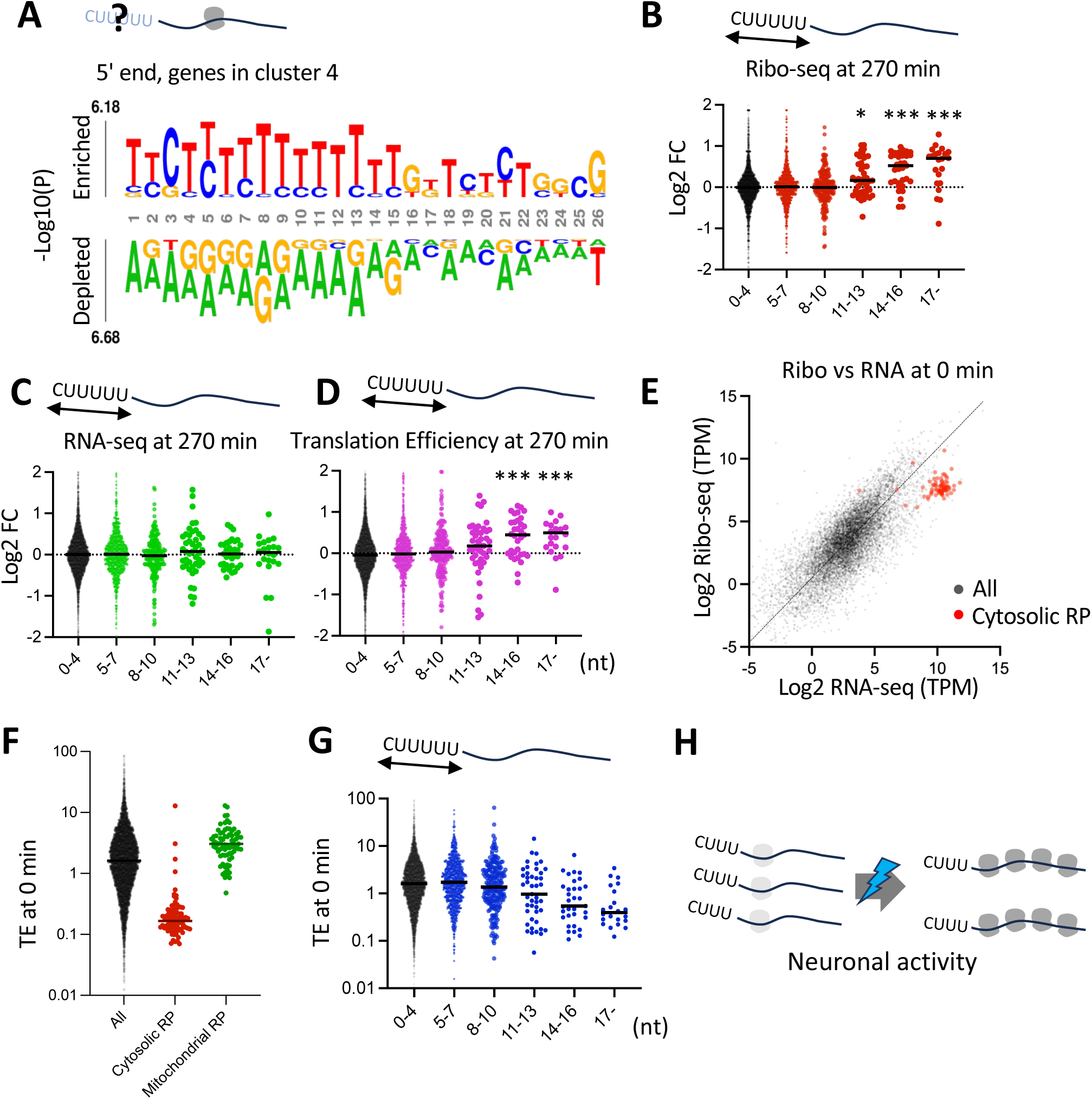
Neuronal activity enhances translation of TOP mRNAs that are basally suppressed. **(A)** Motif-enrichment analysis using kpLogo ^33^, near the 5′ ends of transcripts in the gene cluster showing delayed translational enhancement (cluster 4 shown in Figure 1B). **(B–D)** Log2 fold changes stratified by 5′ terminal pyrimidine tract length (TOP length bins, nt), in **(B)** Ribo-seq, **(C)** RNA-seq, and **(D)** TE. Genes are classified according to oligopyrimidine tract length, allowing up to one purine insertion. *: *P* < 0.05, ***: *P* < 0.001, Dunn’s multiple-comparison test against the “0-4” group. **(E)** Scatter plot comparing baseline mRNA abundance and ribosome occupancy before stimulation. Cytosolic RP mRNAs, highlighted in red, show disproportionately low ribosome occupancy relative to RNA abundance. The dotted line indicates the Deming regression line. **(F)** Baseline TE for all genes, cytosolic RP genes, and mitochondrial RP genes, showing selective translational suppression of cytosolic RP mRNAs. **(G)** Baseline TE as a function of TOP length (nt), showing lower TE in transcripts with longer pyrimidine tracts. **(H)** Model summarizing activity-dependent derepression of TOP mRNA translation in neurons.

Given that longer 5’ TOP tracts confer stronger translational responses to cellular perturbation in other systems ^34^, we examined activity-dependent translational increase as a function of TOP length (Figure 3B-3D). Transcripts with more than 10 consecutive pyrimidines at the 5’ end exhibited robust activity-dependent translational induction without corresponding changes in transcript abundance (Figures 3B-3D and S8). Comparison of Ribo-seq and RNA-seq further showed that TOP mRNAs, especially those encoding cytosolic RPs, are translationally suppressed under basal conditions in neurons (Figure 3E-3G). These results suggest that neuronal activity derepresses TOP mRNA translation (Figure 3H).

### TOP mRNAs are released from RNP granules upon neuronal stimulation

To further examine the cellular mechanism underlying TOP mRNA regulation *in vivo*, we analyzed subcellular localization and activity-dependent dynamics using single-molecule fluorescence *in situ* hybridization (smFISH). Previous work showed that TOP motif-containing mRNAs can localize to RNP granules under cellular stress ^35^. Remarkably, *RpS6* mRNA, which contains a TOP motif, formed prominent puncta in neuronal cell bodies in the basal state (Figure 4A). In contrast, smFISH with control probes against transgenic *UAS-GFP* mRNA ^36^ or total poly(A)-tailed mRNAs produced more diffuse signals (Figure 4A).

**Figure 4.**
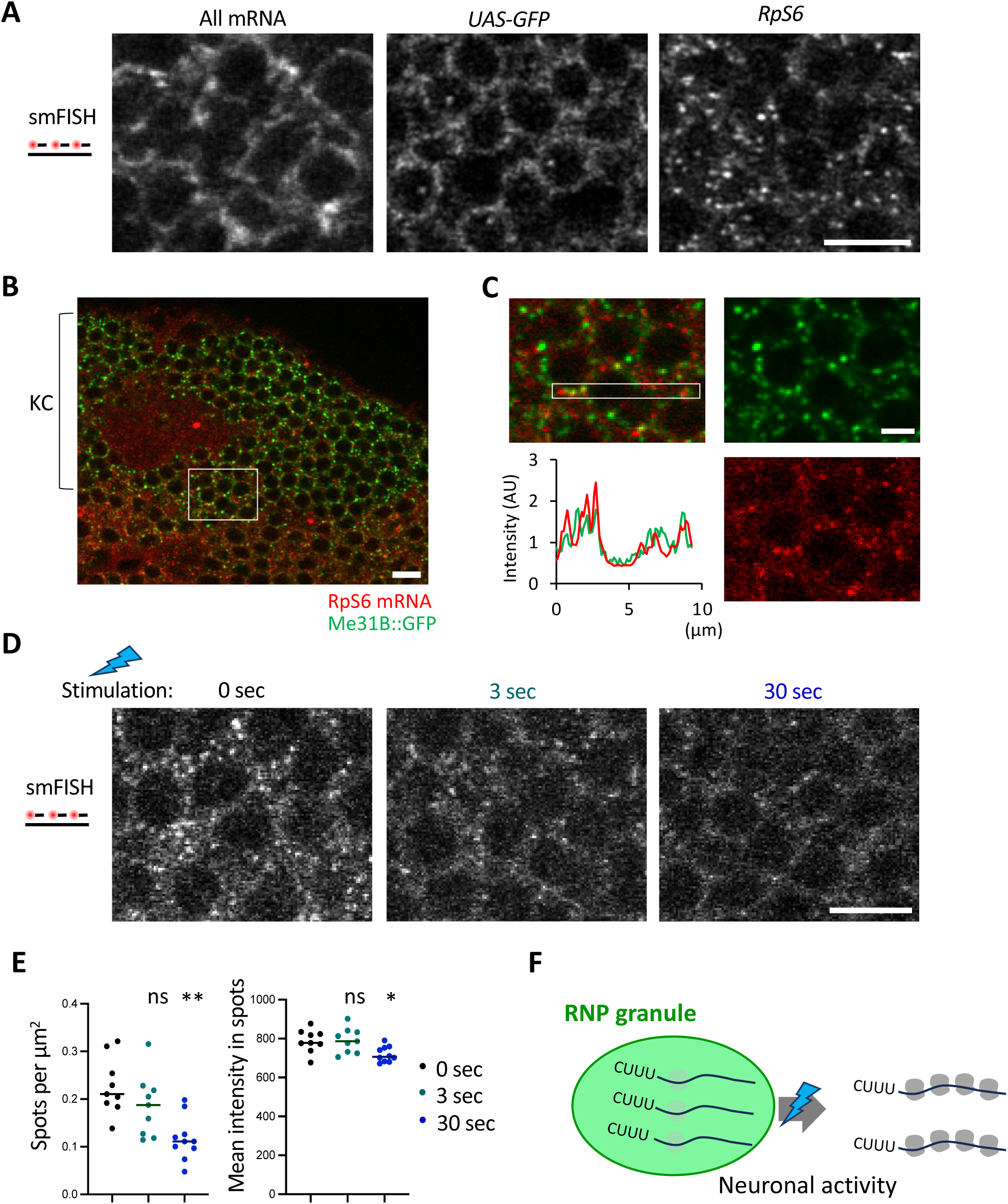
Neuronal activity releases TOP mRNA from RNP granules. **(A)** smFISH images showing distribution of control mRNAs (oligo[dT] probes for total poly[A]-tailed mRNAs and probes specific for GFP mRNA) and RpS6 mRNA in KC somata. RpS6 mRNA forms prominent puncta, whereas control probes show more diffuse signals. Scale bar, 5 µm. **(B)** Representative confocal image of a posterior brain section containing KCs. Green indicates GFP tagged to endogenous Me31B; red indicates *RpS6* mRNA. Scale bar, 5 µm. **(C)** Magnified view of the boxed region in (B), showing colocalization of RpS6 mRNA puncta with Me31B-positive granules. The graph shows a box-scan intensity profile across the region indicated by the rectangle. Scale bar, 2 µm. **(D)** Representative smFISH images of RpS6 mRNA 30 min after 0-, 3-, 30-s optogenetic stimulation, showing activity-induced dispersion of RpS6 mRNA granules. *ChR2XXL* expression was induced after the eclosion upon RU486 feeding with *MB-geneswitch*. Scale bar, 5 µm. **(E)** Quantification of RpS6 mRNA puncta number (spots per µm²) and mean spot intensity in brains stimulated after the indicated duration. Each dot represents one brain. ns, *P* > 0.05, *; *P* < 0.05, **: *P* < 0.01, Dunn’s multiple-comparison test versus control. **(F)** Model illustrating activity-dependent translational release of TOP mRNAs from RNP granules and translational derepression.

To test whether *RpS6* mRNA puncta represent sequestration within translationally inactive RNP granules, we examined colocalization with GFP-tagged endogenous Me31B proteins ^14^. Me31B is the *Drosophila* homolog of DDX6 and a core component of translationally inactive RNP granules, such as P-bodies ^13,37–39^. Under resting conditions, many *RpS6* mRNA puncta colocalized with Me31B-positive granules (Figure 4B and 4C), supporting storage of TOP mRNAs in translationally repressed compartments. To quantify activity-dependent changes in TOP mRNA granules, we expressed *ChR2XXL* in KCs and activated them (3 or 30 seconds). Optogenetic stimulation significantly reduced *RpS6* mRNA granules in KCs (Figure 4D and 4E), while Me31B granules were not strongly affected (Figure S9), suggesting selective release of TOP mRNA from RNP granules. These results demonstrate that neuronal TOP mRNAs are stored in RNP granules under basal conditions and disperse upon neuronal activity to support translation (Figure 4F).

### Larp controls TOP mRNA recruitment into RNP granules and gates long-term memory formation

We next examined how assembly and disassembly of TOP mRNA granules are regulated. In mammalian cells, LARP1 and 4E-BP are known to associate with TOP mRNAs and regulate their translation ^34,40–43^. To assess their roles in activity-dependent translation, we silenced expression of *larp* and *Thor*, the *Drosophila* homologs of *LARP1* and *4E-BP*, in KCs by corresponding transgenic RNAi and examined *RpS6* mRNA granules. Adult-specific knockdown of *larp* significantly reduced *RpS6* mRNA puncta, whereas *Thor* knockdown had only a modest effect (Figure 5A and 5B). Consistently, *larp* knockdown increased basal RpS6 protein levels in KCs, whereas *Thor* knockdown did not (Figure 5C and 5D). Critically, we found that *larp* knockdown occluded the activity-dependent increase in RpS6 protein (Figure 5E and 5F). These results establish Larp as a central regulator of both spatial sequestration and activity-dependent translation of TOP mRNAs.

**Figure 5.**
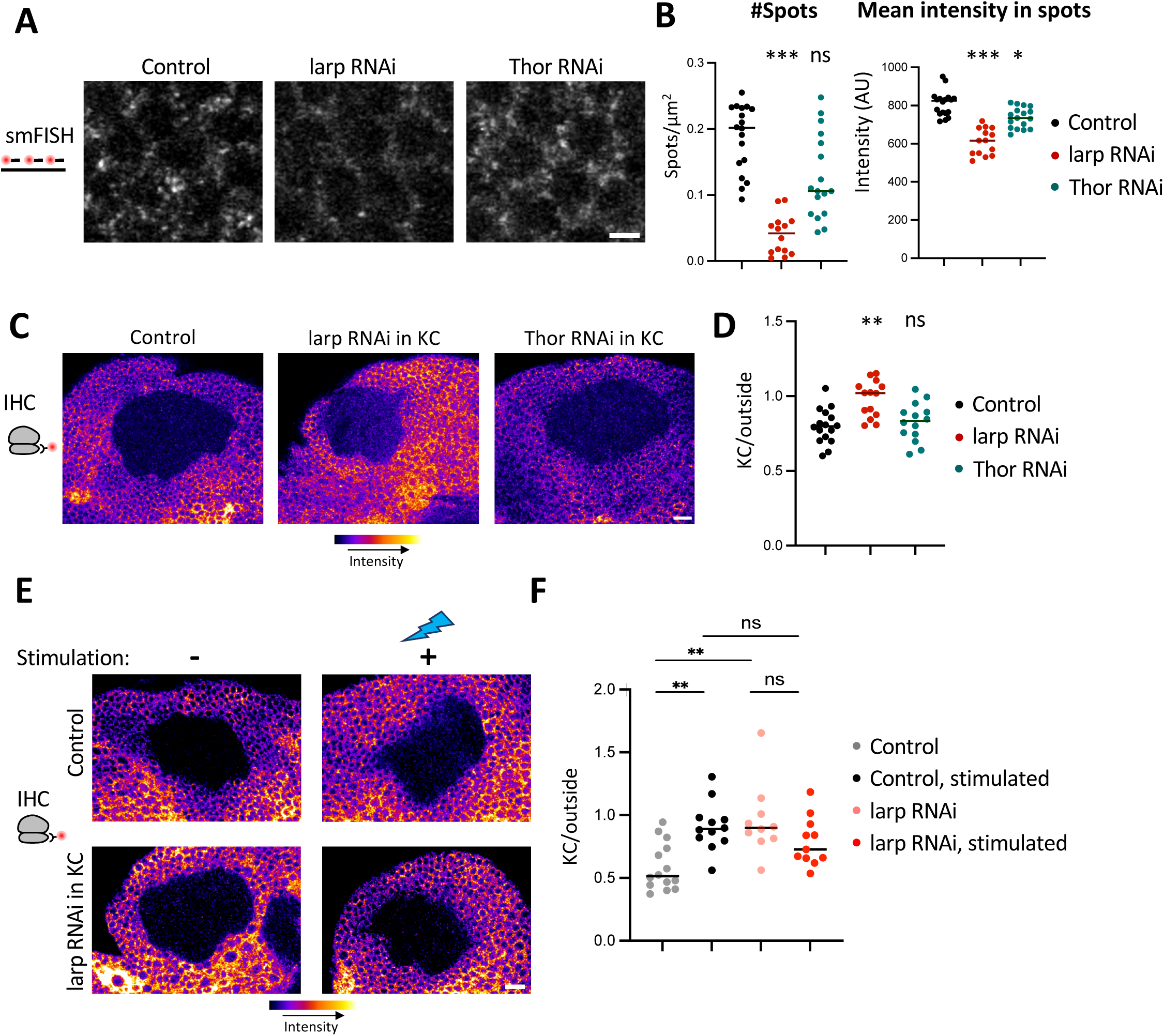
Larp controls RpS6 mRNA granules and activity-dependent RpS6 synthesis. **(A)** Representative smFISH images of *RpS6* mRNA in KCs from control, *larp* RNAi, and *Thor* RNAi, driven by *R13F02-GAL4*. RNAi was induced after eclosion using *tubP-GAL80^ts^*. Scale bar, 2 µm. **(B)** Quantification of *RpS6* mRNA puncta number (spots per µm²) and mean spot intensity across genotypes shown in (A). Each dot represents one brain. ns, *P* > 0.05, *; *P* < 0.05, ***: *P* < 0.001, Dunn’s multiple-comparison test versus control. **(C)** Representative immunohistochemistry images of RpS6 protein in KCs from control, *larp* RNAi, and *Thor* RNAi, driven by *R13F02-GAL4*. RNAi was induced after eclosion using *tubP-GAL80^ts^*. Scale bar, 2 µm. **(D)** Quantification of RpS6 protein signal (KC/outside ratio) across genotypes shown in (C). ns, *P* > 0.05, *; *P* < 0.05, **: *P* < 0.01, Dunn’s multiple-comparison test versus control. **(E)** Representative RpS6 protein images before and 90 min after optogenetic stimulation in control and *larp* RNAi conditions, driven by *R13F02-GAL4*. RNAi was induced after eclosion using *tubP-GAL80^ts^*. Scale bar, 10 µm. **(F)** Quantification of RpS6 protein signal in control and *larp* RNAi animals with and without stimulation. Each dot represents one brain. ns, *P* > 0.05, *; *P* < 0.05, **: *P* < 0.01, Dunn’s multiple-comparison test versus control.

To assess the behavioral significance of this regulation, we examined appetitive olfactory learning. A paired presentation of an odor and sucrose reward induces long-term memory (LTM), which requires *de novo* protein synthesis ^44^. In contrast, non-nutritive sweeteners such as arabinose support only short-term memory (STM; Figure 6A) ^45,46^. Neither *larp* nor *Thor* knockdown in KCs impaired LTM formation after sucrose conditioning (Figure 6B). In contrast, *larp* knockdown enabled significant 24-hr memory formation after arabinose conditioning while leaving STM unchanged (Figure 6C and 6D). These findings identify Larp as a gatekeeper that limits LTM formation after weak reinforcement (Figure 6E).

**Figure 6.**
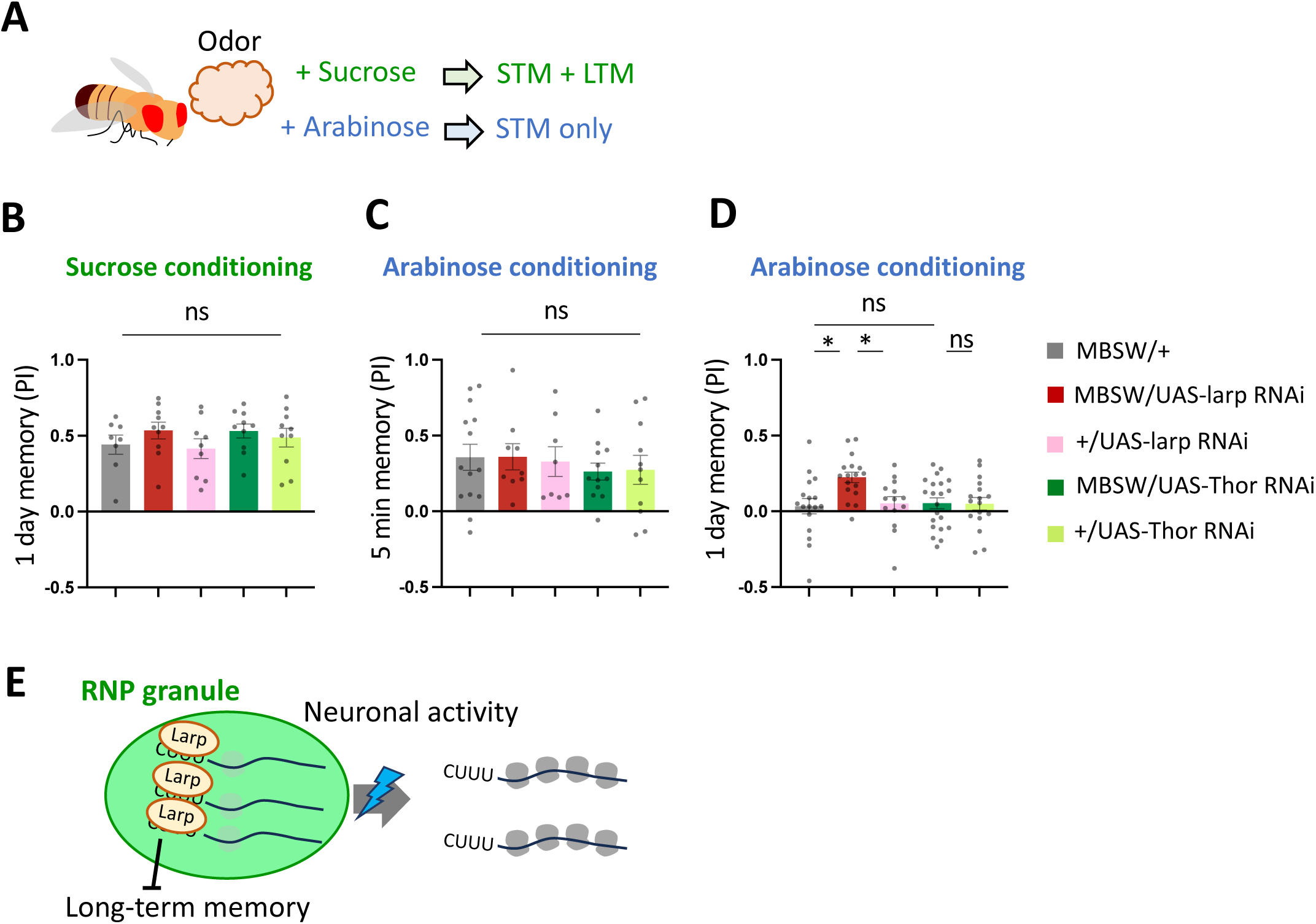
*larp* knockdown in KCs enables long-term memory formation after arabinose conditioning. **(A)** Schematic of appetitive olfactory conditioning. Pairing an odor with sucrose induces long-term memory (LTM), whereas pairing with arabinose typically induces only short-term memory (STM). **(B)** LTM performance index (PI), measured one day after sucrose conditioning, in *larp* or *Thor* knockdown and the control genotypes. Knockdown was performed during the adult by RU486 feeding for 3 days with *MB-switch*. Bars and error bars represent mean and standard error of the mean. ns, *P* > 0.05, Dunn’s multiple-comparison test. **(C)** STM PI, measured 5 min after arabinose conditioning. ns, *P* > 0.05, Dunn’s multiple-comparison test. **(D)** LTM PI, measured one day after arabinose conditioning. ns, *P* > 0.05, *; *P* < 0.05, Dunn’s multiple-comparison test. **(E)** Model illustrating Larp-dependent sequestration of TOP mRNAs in RNP granules as a gate for LTM formation.

### Selective TOP enrichment in ribosomal protein mRNAs is specific to the animal kingdom

Our data demonstrate that TOP mRNAs undergo activity-dependent translational derepression through Larp. While translational regulation through these *cis*- and *trans*-elements is evolutionarily ancient, TOP motifs are not universally enriched in RP mRNAs across species ^47,48^. We thus asked when the association between TOP motifs and RP mRNAs emerged by analyzing the pyrimidine tracts flanking experimentally determined transcription start sites (TSS) across eukaryotic lineages (Figure 7A) ^49–51^. We quantified the local frequency of oligopyrimidine tracts—defined as ten consecutive pyrimidines allowing one purine interruption—within a ±50-nucleotide window centered on each TSS. All tested metazoans (human, mouse, fly, and sea anemone) showed pronounced oligopyrimidine enrichment around TSSs specifically for RP genes (Figure 7B). In stark contrast, this motif was not prevalent among RP genes in fungi or plants (red bread mold, budding yeast, thale cress, and rice; Figure 7B). Notably, the unicellular holozoan *Capsaspora*, a close metazoan relative, showed oligopyrimidine enrichment around TSSs not only in RP genes but broadly across genes (Figure 7B). These results raise the possibility that this regulatory architecture was co-opted during animal evolution to support neuronal control of translational capacity (Figure 7C).

**Figure 7.**
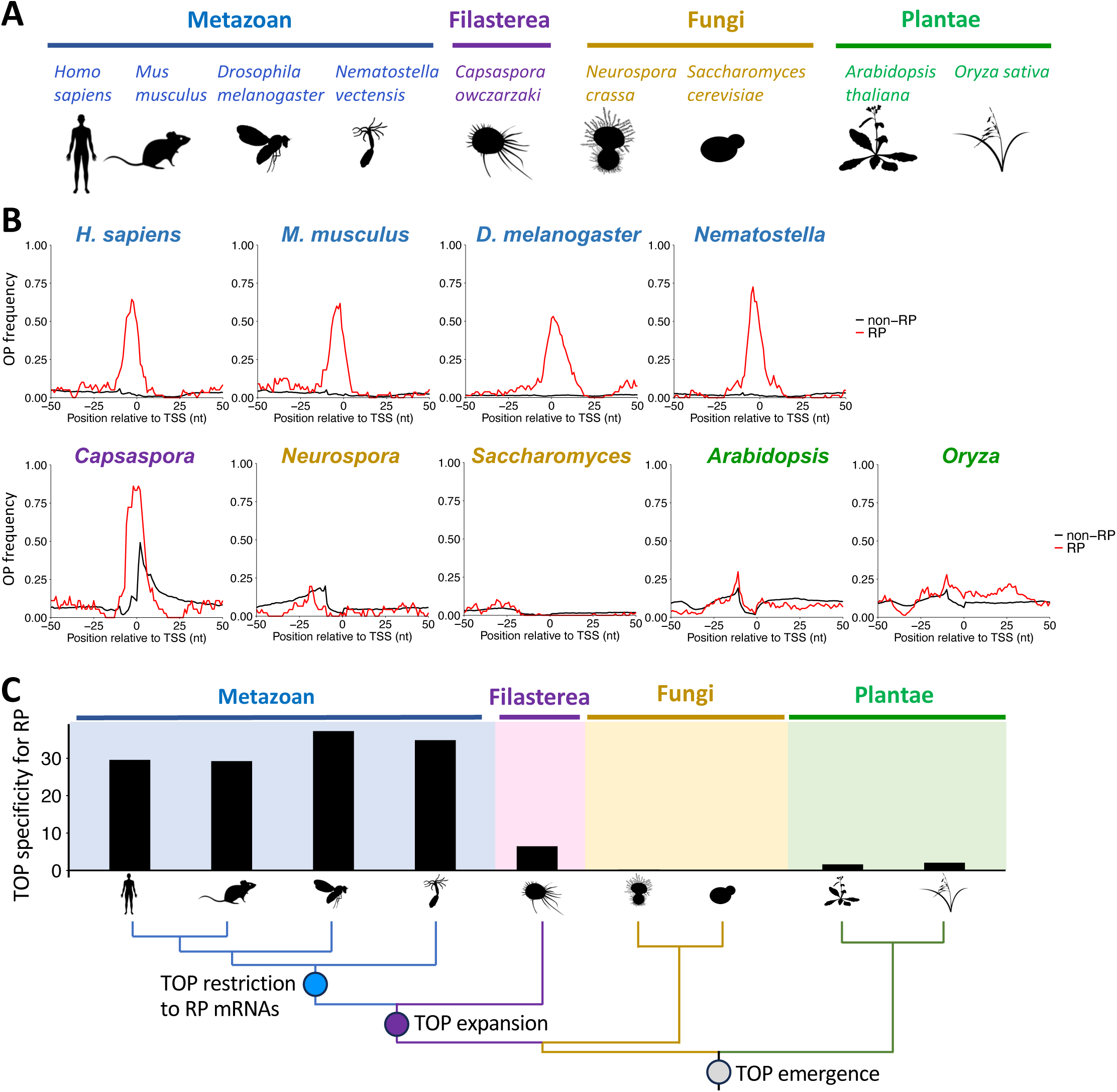
Selective TOP enrichment in ribosomal protein mRNAs is a metazoan feature. **(A)** Schematic showing the species analyzed. **(B)** Frequency of oligopyrimidine tracts around experimentally defined TSSs ^49–51^ for RP genes (red) versus non-RP genes (black) in each species. Oligopyrimidine tracts are defined as ten consecutive pyrimidine nucleotides, allowing up to one purine interruption. Each nucleotide position is scored by tract overlap across the full tract length. **(C)** Quantification of TOP enrichment scores across species, defined as oligopyrimidine frequency in RP genes divided by that in non-RP genes (−3 to 0 nt, relative to the TSSs). A phylogenetic tree with a proposed model of TOP evolution is shown below.

## Discussion

### Hierarchical organization of activity-dependent translational programs

Our study reveals that neuronal activity drives hierarchically organized translational responses in the brain. Optogenetic stimulation of *Drosophila* neurons engages temporally distinct programs that target functionally coherent gene groups. The immediate transient program prioritizes synaptic remodeling factors and neuropeptides (Figure 1), followed by the delayed and dominant program that amplifies the core protein synthesis machinery (Figure 2). This ordered deployment suggests a strategy in which neurons first increase proteins that modify synaptic and circuit properties, then enhance translational capacity to sustain later phases of plasticity.

A key insight from these findings is the functional dissociation between transcriptional and translational responses to neuronal activity. Many immediate early genes transcribed after depolarization encode transcription factors that initiate subsequent gene-expression programs ^1,2,4^. In contrast, the major targets identified here—RPs and translational factors—exhibit minimal changes in corresponding mRNA abundance, indicating regulation primarily at the level of translation (Figures 2, 3, and S3). Given that ribosome biosynthesis is a large metabolic investment ^52^, translational capacity has traditionally been considered most dynamic in contexts such as cell proliferation, growth, and differentiation ^53,54^. In postmitotic cells, ribosome biosynthesis has been reported to be translationally suppressed ^26,29,55,56^. Our data, on the contrary, demonstrate that postmitotic neurons dynamically tune their protein synthesis capacity, consistent with reports of activity-dependent rRNA synthesis and increased ribosome abundance after learning ^57–59^. Together, these findings support a division of labor in which transcription biases plasticity programs, whereas translation acts as a gain-control layer by adjusting the availability of protein synthesis machinery.

### Larp-dependent control of translational capacity couples memory consolidation to energetic investment

Mechanistically, our data suggest that TOP mRNAs are stored within RNP granules and translationally suppressed in a Larp-dependent manner under basal conditions (Figure 4). This is consistent with previous studies reporting stress-dependent sequestration of TOP mRNAs by mammalian LARP1 in RNP granules, including P-bodies and stress granules ^35^. Phosphorylation of LARP1 can interfere with the interaction between LARP1 and TOP mRNAs ^43,60,61^. Larp-dependent storage of TOP mRNAs in neurons (Figure 5) may thus represent a coordinated and economical system that holds ribosome biosynthesis in an idle state while enabling rapid and spatially constrained activation in response to neuronal activity.

This Larp-dependent control of translational capacity aligns with the behavioral effects in LTM formation (Figure 6). LARP1 is known to control nutrition-dependent translation *in vitro* ^62^, and non-nutritive sugars normally support STM but not LTM ^45,46^. Because *de novo* protein synthesis required for LTM is energetically costly, gating translation of TOP mRNAs may prevent costly memory formation when reinforcement is weak ^52,63–68^. Similar to memory enhancement caused by boosting translation ^21^, adult-specific *larp* knockdown in KCs allowed normally short-lasting arabinose memory to persist for 24 hr (Figure 6), suggesting Larp as a gatekeeper that conditionally permits LTM induction. Controlled ribosome production through TOP mRNA may therefore provide a mechanism that couples memory consolidation to both neuronal plasticity and energetic availability.

### Evolutionary emergence of TOP-mediated translational control in metazoans

While TOP motifs and their Larp-dependent regulation seem evolutionarily ancient ^47^, our cross-species TSS analysis suggests that coupling this motif to the RP gene family is a metazoan innovation (Figure 7). Plants and fungi lack this coupling and appear to regulate ribosome biosynthesis primarily through alternative mechanisms, including nutrient-responsive transcriptional control in yeast ^69^. The unicellular holozoan *Capsaspora owczarzaki* may represent an evolutionary intermediate, with oligopyrimidine enrichment near TSS across many genes rather than specifically in RP genes (Figure 7). This lack of RP specificity suggests that TOP motifs may have originally functioned as broad regulatory features that later became selectively associated with RP genes in metazoans. Such specialization may have enabled precise control of translational capacity in nervous systems, where activity-dependent, rapid and scalable cellular responses are required for behavioral adaptation.

## Acknowledgements

We thank Mari Mito (RIKEN) and Genbu Abe (Tottori University) for technical advice on Ribo-seq and *in situ* hybridization. We also thank Dr Akira Nakamura (Kumamoto Univ) and the Bloomington *Drosophila* Stock Center for providing transgenic flies used in this study. Some experiments were performed in the FRIS CoRE, a shared research environment in FRIS, Tohoku University. Computation was supported by the HOKUSAI SailingShip supercomputer facility at RIKEN. This work was supported by JSPS/MEXT (20H05525, 22H05481, 22KK0106, 24H01217, and 25H00986 to H.T.; 21K06369, 21H05713, and 25H02436 to T.I., JP24H02307 to S.I.), JST/FOREST (JPMJFR242Y to TI), Takeda Life Science Research Grant (to TI), the Uehara Memorial Foundation Grant (to TI), RIKEN-Tohoku Univ Science & Technology Hub Collaborative Research Program (to TI and YS) and RIKEN (Pioneering Project to S.I. and Y.S.).

## Declaration of generative AI and AI-assisted technologies in the writing process

During manuscript preparation, the authors used generative AI for language editing and clarity. The authors reviewed, edited, and verified the content and take full responsibility for the final manuscript.

## STAR Methods

### KEY RESOURCES TABLE

| REAGENT or RESOURCE | SOURCE | IDENTIFIER |
| --- | --- | --- |
| **Antibodies** |  |  |
| Anti-FLAG M2 antibody | Sigma-Aldrich | Cat# F1804 |
| Mouse monoclonal anti-RpS6 (clone 54D2) | Cell Signaling Technology | Cat# 54D2 |
| Goat anti-mouse IgG, Cy3 conjugate | Jackson ImmunoResearch | Cat# 155-166-003 |
| **Chemicals, peptides, and recombinant proteins** |  |  |
| Chloramphenicol | Wako Pure Chemical Industries | 030-19452 |
| Cycloheximide | Sigma-Aldrich | C4859-1ML |
| Turbo DNase | Thermo Fisher Scientific | Cat# AM2238 |
| Dynabeads M-280 | Invitrogen | Cat# 11201D |
| RNase I | Epicentre | Cat# N6901K |
| MicroSpin S-400 HR columns | GE Healthcare | Cat# 27-5140-01 |
| Direct-Zol Microprep Kit | Zymo Research | Cat# R2062 |
| SUPERase In | Thermo Fisher Scientific | Cat# AM2694 |
| TRIzol LS | Thermo Fisher Scientific | Cat# 10296010 |
| GlycoBlue | Thermo Fisher Scientific | Cat# AM9515 |
| SuperSep RNA, 15% | Wako Pure Chemical Industries | Cat# 182-02571 |
| T4 polynucleotide kinase | New England Biolabs | Cat# M0201S |
| T4 RNA Ligase 2, truncated KQ | New England Biolabs | Cat# M0373L |
| 5' Deadenylase | New England Biolabs | Cat# M0331S |
| Lambda Exonuclease | New England Biolabs | Cat# M0262L |
| Oligo Clean & Concentrator | Zymo Research | D4060 |
| ProtoScript II | New England Biolabs | Cat# M0368L |
| CircLigaseII ssDNA ligase | Epicentre | Cat# CL9025K |
| NEBNext Poly(A) mRNA Magnetic Isolation Module | New England Biolabs | Cat# E7760 |
| MGIEasy RNA Directional Library Prep kit | MGI tech | Cat# 10000006386 |
| All-trans-retinal | Cayman chemical | Cat# 18449 |
| RU486 (mifepristone) | Tokyo Chemical Industry | Cat# M1732 |
| Hi-Di Formamide | Thermo Fisher Scientific | Cat# 4311320 |
| 3-Octanol | Merck | Cat# 218405-50G |
| 4-Methylcyclohexanol | Sigma-Aldrich | Cat# 153095-250ML |
| Sucrose | Sigma-Aldrich | Cat# S9378-500G |
| D-Arabinose | Tokyo Chemical Industry | Cat# A0513 |
| **Experimental models: Organisms/strains** |  |  |
| *D. melanogaster: UAS-ChR2XXL* | Dawydow et al., 2014 <sup>28</sup> | N/A |
| *D. melanogaster: UAS-RpL3-FLAG* | Chen et al., 2017 <sup>29</sup>; BDSC | BDSC #77132 |
| *D. melanogaster: GMR57C10-GAL4 (nSyb-GAL4)* | BDSC | BDSC #39171 |
| *D. melanogaster: GMR13F02-GAL4* | BDSC | BDSC #48571 |
| *D. melanogaster: tubP-GAL80ts.10* | BDSC | BDSC #7108 |
| *D. melanogaster: Me31B-GFP* | Formicola et al., 2021 <sup>14</sup> | N/A |
| *D. melanogaster: UAS-Larp1 RNAi* | BDSC | BDSC #42578 |
| *D. melanogaster: UAS-thor RNAi* | BDSC | BDSC #36667 |
| *D. melanogaster: MB-geneswitch* | McGuire et al., 2004 <sup>70</sup>; BDSC | BDSC #81013 |
| *D. melanogaster: MB010B* | Aso et al., 2014 <sup>71</sup>; BDSC | BDSC #68293 |
| **Software and algorithms** |  |  |
| STAR aligner | Dobin et al., 2013 <sup>72</sup> | https://doi.org/10.1093/bioinformatics/bts635 |
| DESeq2 | Love et al., 2014 <sup>30</sup> | https://doi.org/10.1186/s13059-014-0550-8 |
| DAVID | Dennis et al., 2003 <sup>73</sup> | https://doi.org/10.1186/gb-2003-4-9-r60 |
| kpLogo | Wu and Bartel, 2017 <sup>33</sup> | https://doi.org/10.1093/nar/gkx323 |
| GraphPad Prism | GraphPad Software | https://www.graphpad.com |
| ImageJ | US National Institute of Health | https://imagej.net/ij/ |
| TANGO (3D spots segmentation ImageJ plugin) | Ollion et al., 2013 <sup>74</sup> | https://doi.org/10.1093/bioinformatics/btt276 |
| **Other** |  |  |
| Confocal microscope Olympus FV1200 | Olympus | FV1200 |
| Confocal microscope Zeiss LSM800 | Zeiss | LSM800 |

### EXPERIMENTAL MODEL AND SUBJECT DETAIL

#### Fly genetics, crosses, and culture

We utilized the following transgenic strains: *UAS–ChR2XXL* ^28^, *UAS–RpL3–FLAG* (BDSC #77132) ^29^, *GMR57C10-GAL4* (nSyb-GAL4) (BDSC #39171), *GMR13F02-GAL4* (BDSC #48571), *tubP-GAL80ts.10* (BDSC #7108), *Me31B-GFP* ^14^, *UAS-Larp1 RNAi* (BDSC #42578), *UAS-thor RNAi* (BDSC #36667), *MB-geneswitch* (BDSC #81013) ^70^. The flies were maintained on standard cornmeal–molasses medium.

Females of the effector lines (*w^1118^;UAS-ChR2-XXL;UAS-RpL3-FLAG, w^1118^;UAS-ChR2XXL, y^1^w^1118^;UAS-ChR2XXL;UAS-Larp1 RNAi, y^1^w^1118^;UAS-ChR2XXL;UAS-thor RNAi*) were crossed to males of the driver lines (*w^1118^;nSyb-GAL4, w^1118^;MB010B, y^1^w^1118^;;MB-geneswitch, y^1^w^1118^;;Me31B-GFP,MB-geneswitch, w^1118^;tubP-GAL80^ts^;GMR13F02-GAL4*), and F1 flies were used for the experiments.

For the experiments involving ChR2 activation, the flies were raised in a dark carton box at 18 °C until eclosion and transferred to 24 °C to minimize activation. 3-7-days-old flies were fed with food supplemented with all-trans-retinal (0.4 mM) for ChR2 activation for 3 days prior to the experiments. When *GAL80^ts^* was used (Figures 2E-F and 5A-F), the adult flies were kept at 29 °C for 3 days instead of 24 °C to induce the GAL4 activity. When *MB-geneswitch* was used (Figures 4D-E and 6), flies were fed with food supplemented with RU486 (0.2 mM) for 3 days prior to the experiments.

#### Library preparation for ribosome profiling and RNA-seq

Adult flies were subjected to a 10 Hz (10 ms ON and 90 ms OFF) optogenetic stimulation for 3 seconds using a custom-made LED tube, kept in a normal food vial in the darkness, and were sampled at defined time points after stimulation (2, 10, 30, 90, and 270 min). Ribosome profiling and transcriptome library preparation were performed in a similar manner to our previous studies ^26,75^ with minor modifications. Briefly, heads were collected by rapid flash freezing and separated from bodies by vortexing and sieving ^76^. Approximately 800 frozen heads were homogenized with 800 µL of frozen droplets of lysis buffer (20 mM Tris-HCl, pH 7.5, 150 mM NaCl, 5 mM MgCl2, 1 mM dithiothreitol, 1% Triton X-100, 100 μg/ml chloramphenicol, and 100 μg/ml cycloheximide). Ribosomes from the neuronal cells were isolated via immunoprecipitation of RpL3–FLAG complexes using anti-FLAG M2 antibody (F1804, Sigma-Aldrich) and Dynabeads M-280 (11201D, Invitrogen). The purified lysate was digested with RNase I (0.005 U/µL, N6901K, Epicentre) at 25 °C for 45 minutes. After isolation of monosomes through MicroSpin S-400 HR columns (27-5140-01, GE Healthcare), footprints were size-selected by denaturing PAGE (17-34 nt) and converted into sequencing libraries in the same manner as in Ichinose et al., 2024 ^26^. The libraries were sequenced with HiSeq4000 or HiSeqX_Ten (Illumina).

For transcriptome analysis, the lysate was prepared as described above, but without RNase digestion and S-400 HR columns. RNA was purified using the Direct-Zol Microprep Kit (R2062, Zymo Research). The libraries were constructed in Azenta Japan Corporation, using the NEBNext Poly(A) mRNA Magnetic Isolation Module (E7760, New England Biolabs) and MGIEasy RNA Directional Library Prep kit (10000006386, MGI tech). The libraries were sequenced with DNB-seq with an option of paired-end reads for 150 bases.

#### Data analysis

Sequencing data processing and quantification were performed in a similar manner to Ichinose et al., 2024 ^26^ with minor modifications. Briefly, adapters were trimmed, low-quality bases were removed, and reads mapping to noncoding RNAs were filtered. Remaining reads were aligned to the *Drosophila* reference genome (dm6) using STAR ^72^. For ribosome profiling, fragments ranging from 21-36 nt were analyzed, and the position of the P-site was estimated as 12 nt downstream of the 5’ end. RNA-seq reads were quantified per gene using the same annotation framework. Fold change was calculated as mean TPM on CDS (Ribo-seq) or whole transcript (RNA-seq), normalized by TPM of the control samples (no stimulation). At least two biological replicates were analyzed per condition, and the number of unique reads mapped on mRNAs is as follows.

Ribo-seq:

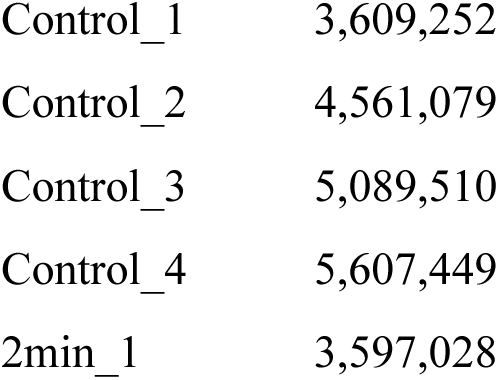
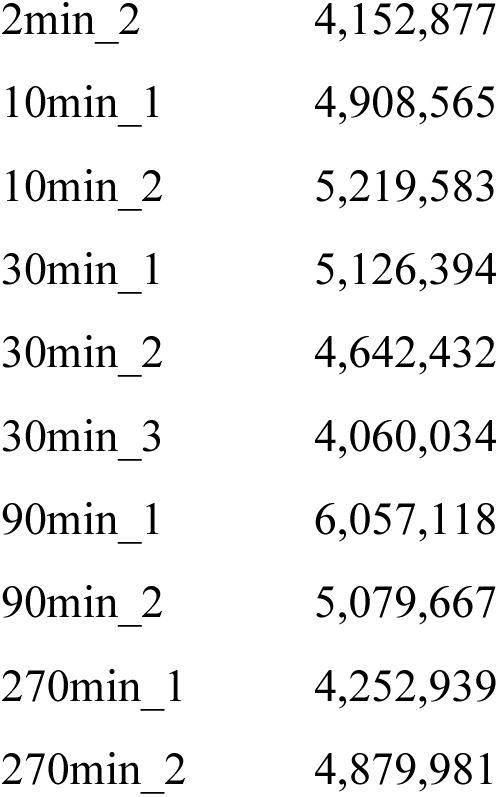

|  |  |
| --- | --- |
| Control_1 | 3,609,252 |
| Control_2 | 4,561,079 |
| Control_3 | 5,089,510 |
| Control_4 | 5,607,449 |
| 2min_1 | 3,597,028 |
| 2min_2 | 4,152,877 |
| 10min_1 | 4,908,565 |
| 10min_2 | 5,219,583 |
| 30min_1 | 5,126,394 |
| 30min_2 | 4,642,432 |
| 30min_3 | 4,060,034 |
| 90min_1 | 6,057,118 |
| 90min_2 | 5,079,667 |
| 270min_1 | 4,252,939 |
| 270min_2 | 4,879,981 |

RNA-seq:

|  |  |
| --- | --- |
| Control_1 | 43,554,787 |
| Control_2 | 52,705,043 |
| Control_3 | 18,877,601 |
| Control_4 | 19,712,382 |
| 30min_1 | 36,241,293 |
| 30min_2 | 38,098,416 |
| 30min_3 | 19,529,435 |
| 90min_1 | 16,920,072 |
| 90min_2 | 22,858,556 |
| 270min_1 | 21,296,010 |
| 270min_2 | 22,002,617 |

Differential translation and transcription were assessed using count-based statistical modeling (DESeq2 ^30^). Translational efficiency (TE) was computed as TPM on the CDS (Ribo-seq) divided by TPM of the transcript (RNA-seq). Heatmaps (Figures 2A and S2) were generated in *R* using the pheatmap package. Genes (rows) were hierarchically clustered using Euclidean distance.

Gene Ontology (GO) term enrichment analysis (Figures 1C, 2C, and S1) was performed on the Database for Annotation, Visualization, and Integrated Discovery (DAVID) ^73^, and was visualized in *R*. The background gene set was defined as all the genes showing at least one read in Ribo-seq in all conditions. Motif-enrichment (Figure 3A) was analyzed using k-mer probability logo ^33^. The 26 bases on the 5’ end, according to the annotation (*Drosophila* genome dm6), were analyzed with all genes as the background.

Statistical tests were performed on GraphPad Prism 10.

### Immunohistochemistry and fluorescent in-situ hybridization

Immunohistochemistry (Figures 2E-F and 5C-F) was performed as previously described with minor modifications ^77^. Briefly, male fly brains were dissected in 2% formaldehyde in PBS, fixed in 4% formaldehyde in PBS for 30 min at room temperature, washed three times with PBST, blocked with 3% goat serum in PBST for 30 min, then incubated with the primary antibody solution at 4°C overnight (mouse anti-RpS6 (1:2000, Cell Signaling, 54D2). Subsequently, the brains were washed three times with PBST, incubated with the secondary antibody solution at 4°C overnight (anti-mouse Cy3 (1:1000; Jackson ImmunoResearch, 155-166-003)), and mounted with 86% glycerol in 1x Tris buffer (20 mM, pH 7.4).

Fluorescent in-situ hybridization (Figures 4A-C, 5A, and S9) was performed as previously described ^26^. Male fly brains were dissected in 2% formaldehyde in PBS, fixed in PBS containing 3% formaldehyde, 1% glyoxal, and 0.1% methanol for 30 min at room temperature, followed by three quick washes with PBST. The buffer was then exchanged to the wash solution (10% Hi-Di formamide (Thermo Fisher Scientific; 4311320) in 2x saline sodium citrate) and was incubated at 37°C for 16 hr. The probes were rinsed three times with preheated wash solution at 37°C, followed by three washes for 10 min at room temperature. The brains were mounted with 86% glycerol in 1x Tris-HCl buffer (20 mM, pH 7.4).

For the experiments involving optogenetic activation of Kenyon cells (Figures 2E-F, 4D-E, S9, and 5E-F), flies were fed with all-trans-retinal (0.4 mM) for 3 days and subjected to a 10 Hz (10ms ON and 90ms OFF) optogenetic stimulation for 30 seconds using a custom-made LED tube. The brains were dissected under red LED and fixed in the darkness to avoid potential ChR2 activation during preparation.

The brains were imaged by confocal microscopy (Olympus FV1200 or Zeiss LSM800) under identical acquisition settings across genotypes and conditions. To quantify the fluorescent intensity (Figure 2F, 5D, and 5F), 5 slices (15 μm square) covering the Kenyon cells or the other regions were cropped per brain, and mean intensity was measured in ImageJ. To quantify mRNA aggregation (Figures 4E and 5B), slices were cropped similarly, and an ImageJ plugin “3D spots segmentation” ^74^ was utilized to count the identified spot number and to measure the intensity inside. These values were averaged across slices, and one value was calculated for each brain.

### Behavioral measurement

Appetitive olfactory learning measurements (Figure 6) were conducted in a similar manner to our previous studies ^78,79^. Flies were wet-starved so that the mortality in the test became around 10%. For conditioning, a group of approximately 50 flies in a tube received alternately 1.2% 3-octanol (Merck) or 2% 4-methylcyclohexanol (Sigma-Aldrich), respectively, for 1 minute in a constant air flow with or without reinforcements. Dried sucrose (1.5 mL of 2 M solution on 74 mm x 42 mm filter paper) or arabinose (1.5 mL of 3 M solution) was presented as a reinforcement. Conditioned flies were kept starved until the test. Memory performance was assessed in a choice test between the two odors, and the performance index was calculated as (#CS+ - #CS-)/(#CS+ + #CS-), where #CS+ and #CS-indicate the number of flies in the associated or control odors, respectively. Performance indices were calculated every second and averaged across 60 sec to 120 sec after the onset of the choice ^79^.

### Sequence analysis around transcription start sites

Cross-species analysis of oligopyrimidine tract enrichment around transcription start sites (TSS) (Figure 7) was performed using publicly available TSS datasets with single-nucleotide or high-resolution definitions. TSS annotations for *Homo sapiens, Mus musculus, Nematostella vectensis, Capsaspora owczarzaki, Neurospora crassa*, and *Oryza sativa* were obtained from ^49^.

For *Drosophila melanogaster*, TSS annotation was obtained from Ensembl Metazoa.

For *Saccharomyces cerevisiae* and *Arabidopsis thaliana*, TSS annotation was derived from cluster-based CAGE-seq datasets, obtained from YeasTSS (http://www.yeastss.org/), and from ^51^, respectively, and dominant peak positions (dominant_ctss for *S. cerevisiae* and TC_peak for *A. thaliana*) were used. When multiple TSSs were annotated for one gene, the TSS closest to the start codon but located ≥10 bp upstream in intergenic regions was selected.

For each annotated protein-coding gene, the DNA sequences flanking the representative TSS were extracted in a ±50 nt window. Oligopyrimidine tracts were defined as ten or more consecutive pyrimidines (C/T) allowing one purine interruption within the tract, and tract frequency was quantified across the TSS window.

---

**Figure S1. Quality control of the neuron-specific Ribo-seq**

**(A)** MA-plot. The present neuron-specific Ribo-seq dataset was compared to the whole-head sample in our previous study ^26^. The x- and y- axes represent the average and fold change between the two datasets. Well-known marker genes, described below, are highlighted. Neuron (green): *nSyb, Ilp2, Lk, Trh, Shaw, nAChRa1, FoxP, Gad1, ChAT, Vglut, para, Sh, Tbh, ppk, AstC, rad, NPF, mAChR-B, elav, Rab3, Syta, Hug*. Glia (blue): *alrm, repo, moody, wrapper, nrv2, zyd, Gat, Balat*. Muscle (brown): *Mhc, wupA, Tm2, sls, up, Mp20, Mlp60A*. Fat body (yellow): *Lsd-2, FASN1, Yp1, Yp2, apolpp, Lsp2, fit, AkhR, fon*. Epidermis (dark blue): *Cpr47Ea, Cpr49Af, Cpr65Au, Cpr67Fb*). Hemocytes (red): *Hml, eater, NimC1, NimC2*. Olfactory support cells (purple): *Obp19b, Obp49a, Obp56g, Obp83ef, Obp99a*.

**(B)** Meta-genome plot of estimated ribosomal P-site distribution around the start and the stop codons. RNA fragments of 32-nt length, which is the most abundant, are analyzed.

**(C)** Comparison of Ribo-seq data with or without expression of *ChR2XXL*. Pearson’s correlation coefficient is calculated.

**Figure S2. All differentially translated transcripts**

Heatmap showing significant up- or down-regulation in Ribo-seq with the gene names. Genes are hierarchically clustered. Left panel: clusters 1–3; right panel: cluster 4. Note that the same heatmap is shown in Figure 1B, but without the gene names.

**Figure S3. GO term analysis at 30 min and 90 min**

GO enrichment analysis of translationally upregulated genes at 30 min (left) and 90 min (right) after optogenetic stimulation. Dot size indicates gene number, and color indicates enrichment significance (−log10(FDR)).

**Figure S4. Time course of translatome and transcriptome changes**

Time-course profiles of selected gene families. Curves show Log2FC in Ribo-seq and RNA-seq measurements. Asterisks indicate *FDR* < 0.05.

**Figure S5. Mitochondrial ribosomal proteins are not translationally enhanced**

Log2FC of mitochondrial ribosomal protein in Ribo-seq, RNA-seq, and translation efficiency (TE).

**Figure S6. Consistent results with a previous activity-dependent transcriptome study**

The top 12 activity-regulated genes in Chen et al., 2016 (eLife) are plotted according to their dataset ^7^ (ChR2-XXL activation, 60 min after stimulation) and our RNA-seq dataset (90 min).

**Figure S7. TOP motif enrichment in ribosomal proteins**

Motif enrichment analysis near the 5’ termini of transcripts encoding cytosolic ribosomal proteins (k-mer probability logo ^33^).

**Figure S8. Fold change of TOP mRNAs in RNA-seq and Ribo-seq**

Scatter plots showing fold changes of TOP-length-binned transcripts in Ribo-seq, RNA-seq, and translation efficiency at 30, 90, and 270 min after stimulation. All the genes are classified according to the oligopyrimidine tract length, allowing up to one purine insertion.

**Figure S9. Me31B puncta are not strongly affected by optogenetic stimulation**

Representative images of Me31B::GFP granules (green) and RpS6 mRNA smFISH signal (red) with or without optogenetic stimulation (30 min after stimulation). *ChR2XXL* was expressed by using *MB-geneswitch*. In contrast to the rapid reduction of RpS6 mRNA puncta (Figure 4D), Me31B-positive granules are not strongly altered under the same stimulation condition.

