## Supplementary figure for "Activity-dependent synthesis of translational machinery through dynamic remodeling of ribonucleoprotein granules"

### Figure S1

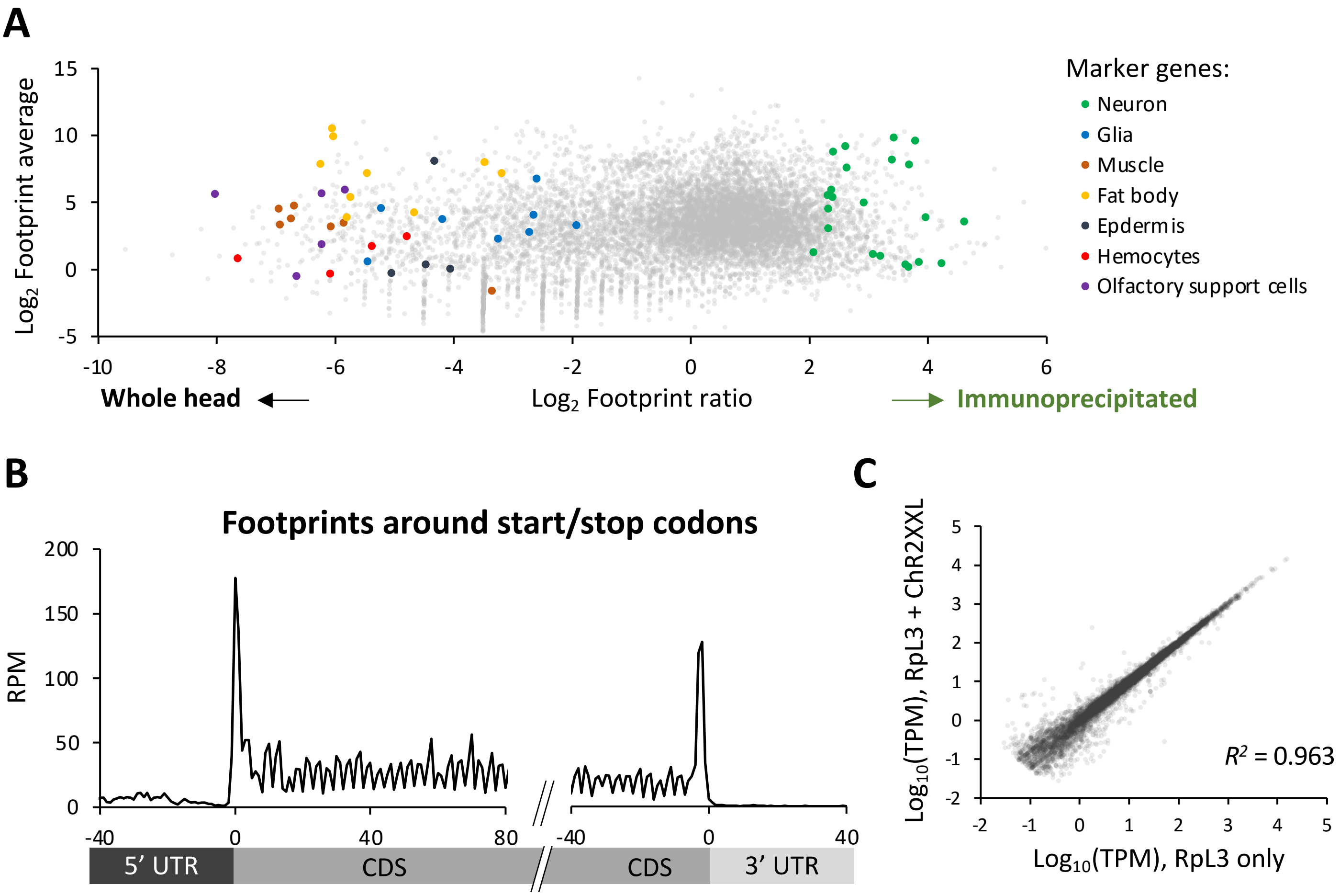
A
15
10
5
0
-5
Log2 Footprint average
-10
-8
-6
-4
-2
0
2
4
6
Whole head
Log2 Footprint ratio
Immunoprecipitated
Marker genes:
Neuron
Glia
Muscle
Fat body
Epdermis
Hemocytes
Olfactory support cells
B
Footprints around start/stop codons
200
150
100
50
0
RPM
-40
0
40
80
-40
0
40
5' UTR
CDS
CDS
3' UTR
C
Log10(TPM), RpL3 + ChR2XXL
5
4
3
2
1
0
-1
-2
-2
-1
0
1
2
3
4
5
R2 = 0.963
Log10(TPM), RpL3 only

### Figure S2

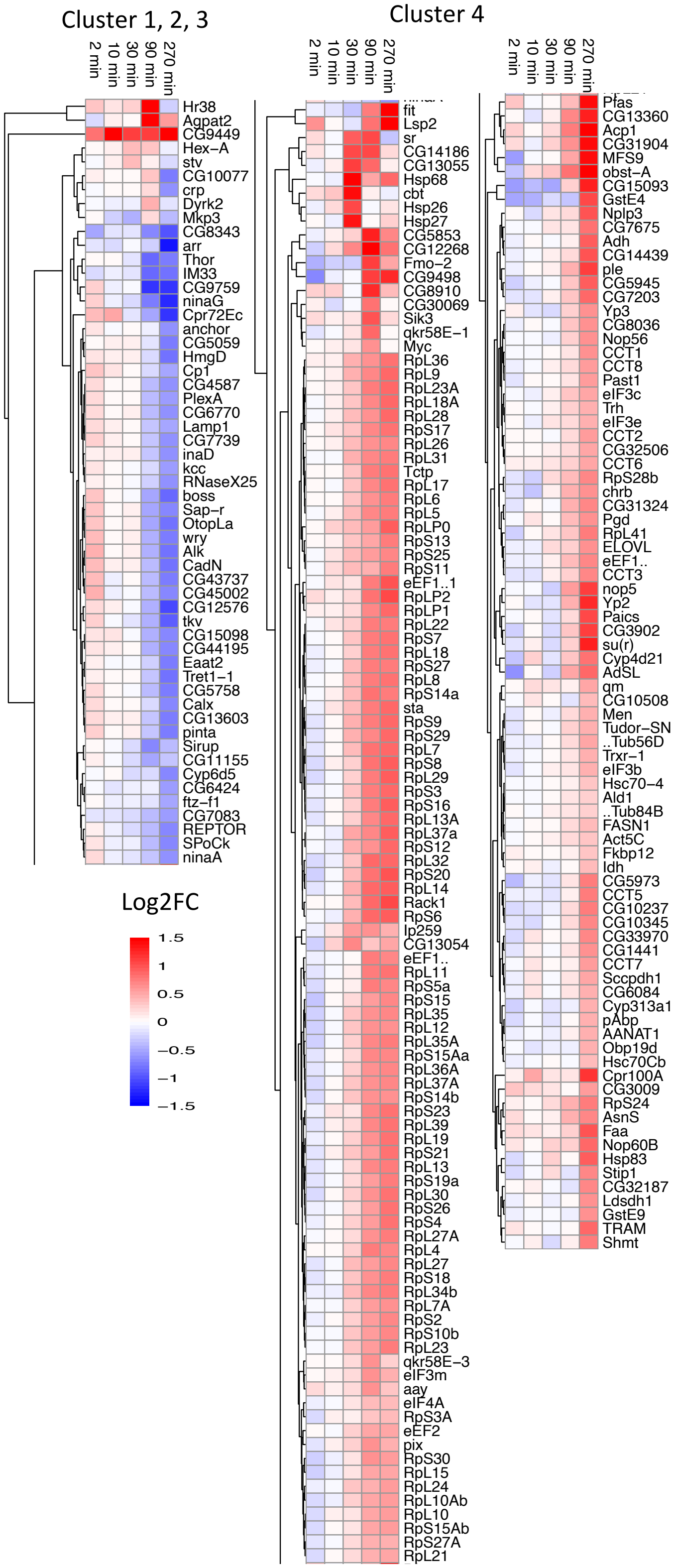

Cluster 1, 2, 3
2 min
10 min
30 min
90 min
270 min
Hr38
Agpat2
CG9449
Hex−A
stv
CG10077
crp
Dyrk2
Mkp3
CG8343
arr
Thor
IM33
CG9759
ninaG
Cpr72Ec
anchor
CG5059
HmgD
Cp1
CG4587
PlexA
CG6770
Lamp1
CG7739
inaD
kcc
RNaseX25
boss
Sap−r
OtopLa
wry
Alk
CadN
CG43737
CG45002
CG12576
tkv
CG15098
CG44195
Eaat2
Tret1−1
CG5758
Calx
CG13603
pinta
Sirup
CG11155
Cyp6d5
CG6424
ftz−f1
CG7083
REPTOR
SPoCk
ninaA
Log2FC
1.5
1
0.5
0
−0.5
−1
−1.5
Cluster 4
2 min
10 min
30 min
90 min
270 min
fit
Lsp2
sr
CG14186
CG13055
Hsp68
cbt
Hsp26
Hsp27
CG5853
CG12268
Fmo−2
CG9498
CG8910
CG30069
Sik3
qkr58E−1
Myc
RpL36
RpL9
RpL23A
RpL18A
RpL28
RpS17
RpL26
RpL31
Tctp
RpL17
RpL6
RpL5
RpLP0
RpS13
RpS25
RpS11
eEF1..1
RpLP2
RpLP1
RpL22
RpS7
RpL18
RpS27
RpL8
RpS14a
sta
RpS9
RpS29
RpL7
RpS8
RpL29
RpS3
RpS16
RpL13A
RpL37a
RpS12
RpL32
RpS20
RpL14
Rack1
RpS6
Ip259
CG13054
eEF1..
RpL11
RpS5a
RpS15
RpL35
RpL12
RpL35A
RpS15Aa
RpL36A
RpL37A
RpS14b
RpS23
RpL39
RpL19
RpS21
RpL13
RpS19a
RpL30
RpS26
RpS4
RpL27A
RpL4
RpL27
RpS18
RpL34b
RpL7A
RpS2
RpS10b
RpL23
qkr58E−3
eIF3m
aay
eIF4A
RpS3A
eEF2
pix
RpS30
RpL15
RpL24
RpL10Ab
RpL10
RpS15Ab
RpS27A
RpL21
2 min
10 min
30 min
90 min
270 min
Plas
CG13360
Acp1
CG31904
MFS9
obst−A
CG15093
GstE4
Nplp3
CG7675
Adh
CG14439
ple
CG5945
CG7203
Yp3
CG8036
Nop56
CCT1
CCT8
Past1
eIF3c
Trh
eIF3e
CCT2
CG32506
CCT6
RpS28b
chrb
CG31324
Pgd
RpL41
ELOVL
eEF1..
CCT3
nop5
Yp2
Paics
CG3902
su(r)
Cyp4d21
AdSL
qm
CG10508
Men
Tudor−SN
..Tub56D
Trxr−1
eIF3b
Hsc70−4
Ald1
..Tub84B
FASN1
Act5C
Fkbp12
Idh
CG5973
CCT5
CG10237
CG10345
CG33970
CG1441
CCT7
Sccpdh1
CG6084
Cyp313a1
pAbp
AANAT1
Obp19d
Hsc70Cb
Cpr100A
CG3009
RpS24
AsnS
Faa
Nop60B
Hsp83
Stip1
CG32187
Ldsdh1
GstE9
TRAM
Shmt

**Figure S3**

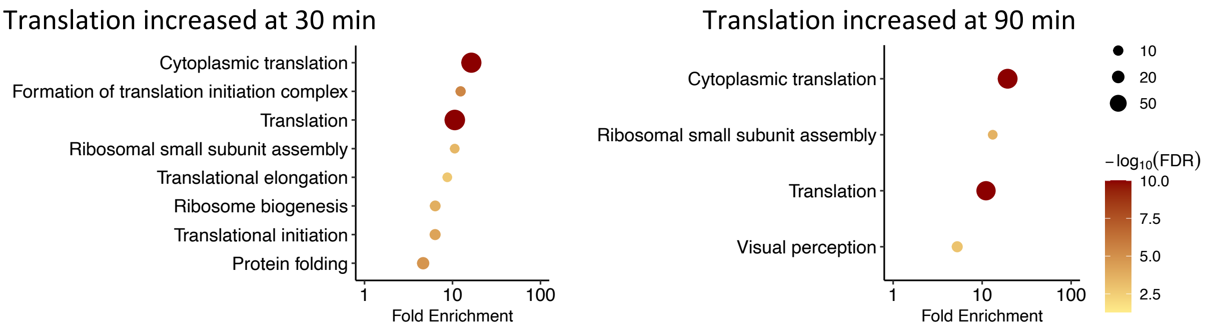
Translation increased at 30 min
Cytoplasmic translation
Formation of translation initiation complex
Translation
Ribosomal small subunit assembly
Translational elongation
Ribosome biogenesis
Translational initiation
Protein folding
1
10
100
Fold Enrichment
Translation increased at 90 min
Cytoplasmic translation
Ribosomal small subunit assembly
Translation
Visual perception
1
10
100
Fold Enrichment
10
20
50
$-\log_{10}(FDR)$
10.0
7.5
5.0
2.5

### Figure S4

#### Ribosome assembly

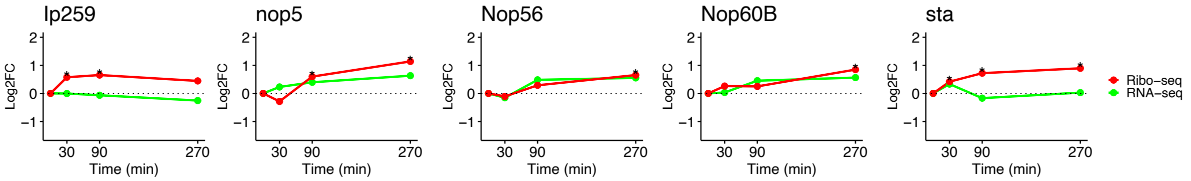


#### Translational factors

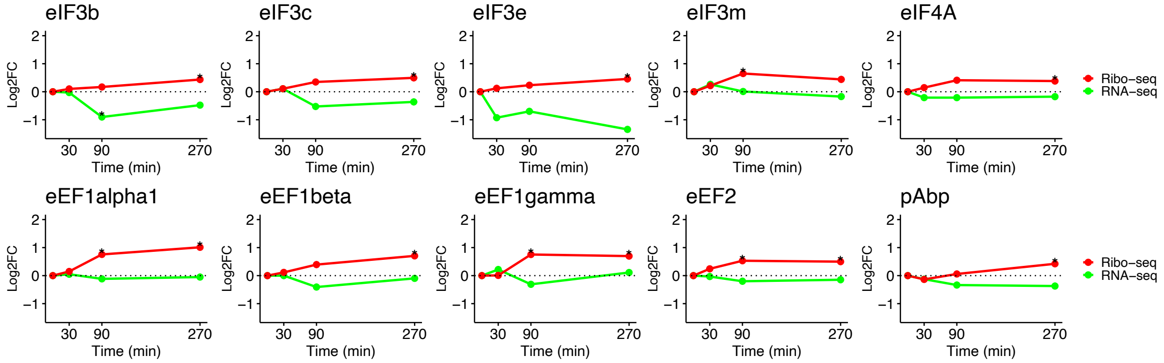


#### Heat shock proteins and chaperonin complex

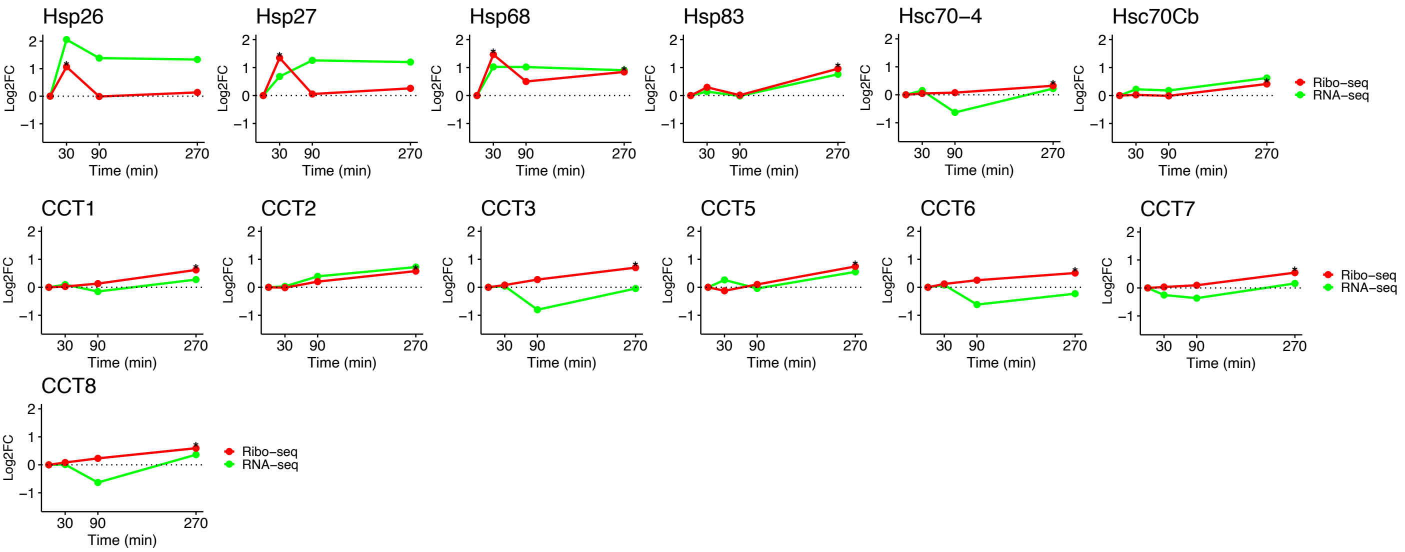


#### Cytoskeleton

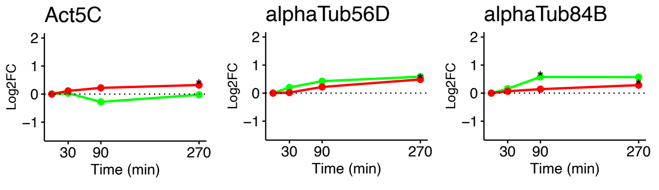


#### Neurotransmitter-related

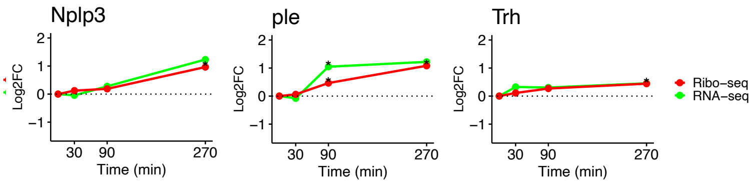


#### Oxidative stress

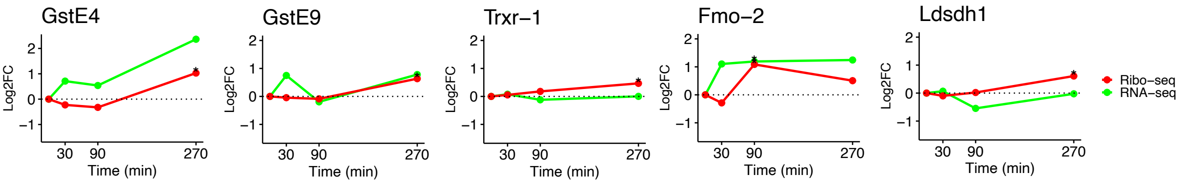

### Figure S5

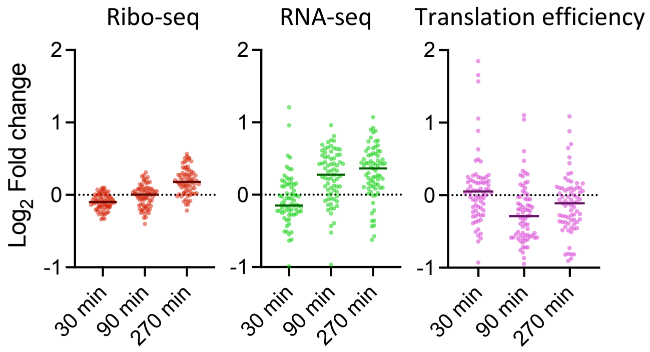
Ribo-seq
RNA-seq
Translation efficiency
Log2 Fold change
2
1
0
-1
30 min
90 min
270 min

### Figure S6

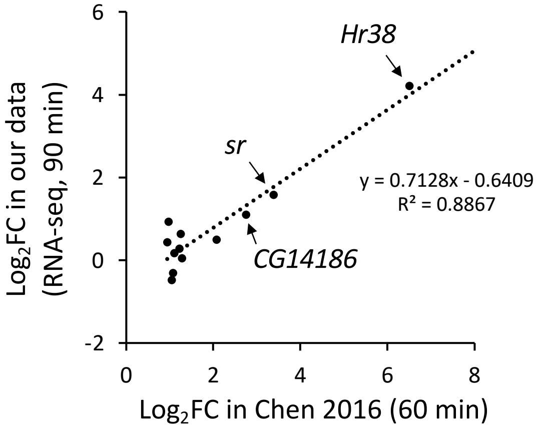

6
4
2
0
-2
Log2FC in our data
(RNA-seq, 90 min)
Hr38
sr
CG14186
y = 0.7128x - 0.6409
R² = 0.8867
0
2
4
6
8
Log2FC in Chen 2016 (60 min)

### Figure S7

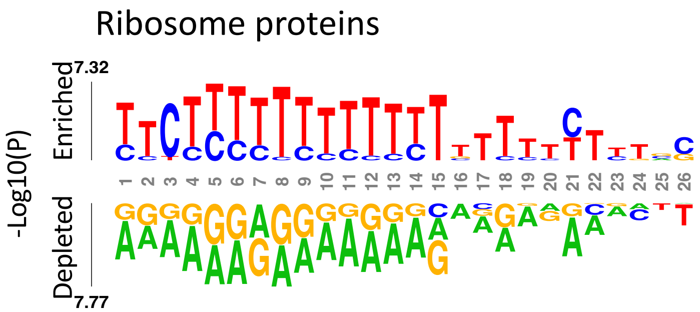
Ribosome proteins
-Log10(P)
Enriched
Depleted
7.32
7.77
1 2 3 4 5 6 7 8 9 10 11 12 13 14 15 16 17 18 19 20 21 22 23 24 25 26

### Figure S8

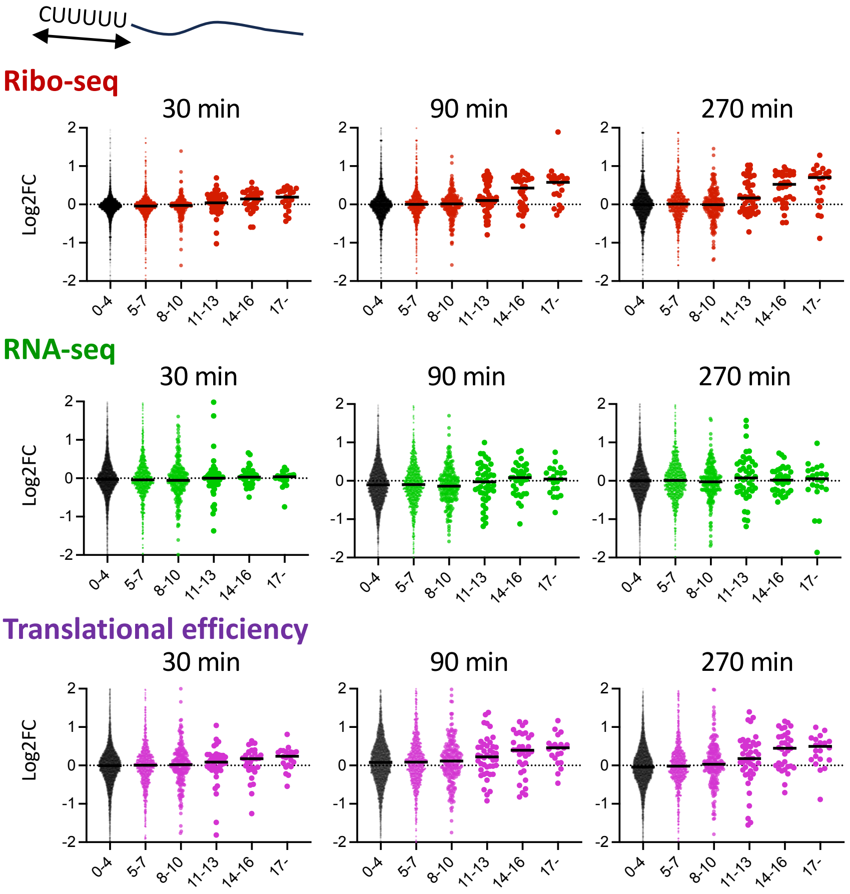
CUUUUU
Ribo-seq
30 min
90 min
270 min
Log2FC
2
1
0
-1
-2
0-4
5-7
8-10
11-13
14-16
17-
RNA-seq
30 min
90 min
270 min
Log2FC
Translational efficiency
30 min
90 min
270 min
Log2FC

### Figure S9

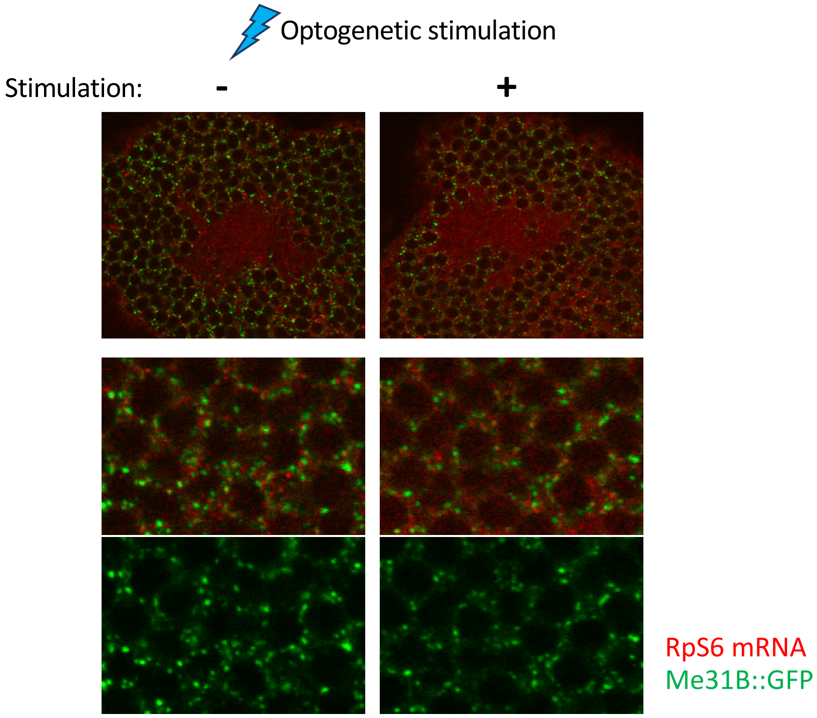
